# SparrKULee: A Speech-evoked Auditory Response Repository of the KU Leuven, containing EEG of 85 participants

**DOI:** 10.1101/2023.07.24.550310

**Authors:** Bernd Accou, Lies Bollens, Marlies Gillis, Wendy Verheijen, Hugo Van hamme, Tom Francart

## Abstract

Researchers investigating the neural mechanisms underlying speech perception often employ electroencephalography (EEG) to record brain activity while participants listen to spoken language. The high temporal resolution of EEG enables the study of neural responses to fast and dynamic speech signals. Previous studies have successfully extracted speech characteristics from EEG data and, conversely, predicted EEG activity from speech features.

Machine learning techniques are generally employed to construct encoding and decoding models, which necessitate a substantial amount of data. We present SparrKULee: A Speech-evoked Auditory Repository of EEG, measured at KU Leuven, comprising 64-channel EEG recordings from 85 young individuals with normal hearing, each of whom listened to 90-150 minutes of natural speech. This dataset is more extensive than any currently available dataset in terms of both the number of participants and the amount of data per participant. It is suitable for training larger machine learning models. We evaluate the dataset using linear and state-of-the-art non-linear models in a speech encoding/decoding and match/mismatch paradigm, providing benchmark scores for future research.

## Background & Summary

In order to study the neural processing of speech, recent studies have presented natural running speech to participants while the electroencephalogram (EEG) was recorded. Currently, regression is used to either decode features from the speech stimulus from the EEG (also known as a backward model)^1–5^, to predict the EEG from the speech stimulus^1,6^ (forward model), or to transform both EEG and speech stimulus to a shared space^7,8^ (hybrid model). Deep neural networks have recently been proposed for auditory decoding and have obtained promising results^4,5,9–12^.

All previously mentioned methods require EEG recordings of the participants with strict time alignment to the speech stimulus. This time alignment is necessary due to the time-locked neural tracking of the speech stimulus at a millisecond scale (e.g., auditory brainstem responses (ABR)), which can last up to 600 ms^13^. As this data is personal and expensive to collect, there is a need for more public datasets that researchers can use to benchmark and train their models.

Table 1 presents an overview of currently available public datasets of EEG recordings of people listening to natural speech. These studies have generated 87.7 hours of EEG data from 133 participants listening to clean speech and speech-in-noise in their native language. However, this amount of data is relatively small compared to datasets in other domains, such as automatic speech recognition, and needs to be increased for training models due to the low signal-to-noise ratio of auditory EEG. Additionally, combining the data from these studies for model training is challenging due to differences in the authors’ signal acquisition equipment, measurement protocols, and preprocessing methods.

**Table 1.** Overview of currently publicly available single-speaker datasets.

| Dataset | Ref | Speech material | Language | Participants | Time per participant (min) | Total time (min) |
| --- | --- | --- | --- | --- | --- | --- |
| Broderick | 14 | clean speech | English | 19 | 60 | 1140 |
|  |  | time-reversed speech |  | 10 | 60 | 600 |
|  |  | speech-in-noise |  | 21 | 30 | 630 |
| DTU Fuglsang | 15 | clean speech | Danish | 18 | 8.3 | 150 |
| Etard | 16 | clean speech | English | 18 | 10 | 180 |
|  |  | speech-in-noise |  | 18 | 30 | 540 |
|  |  | foreign language speech | Dutch | 12 | 40 | 480 |
| Weissbart | 17 | clean speech | English | 13 | 40 | 520 |
| Brennan | 41 | clean speech | English | 49 | 12.4 | 610 |
| Vanheuseden | 42 | clean speech | English | 17 (* mild to severe hearing loss) | 24 | 410 |
| SparrKULee |  | clean speech | Dutch | 85 | 110 | 9320 |
|  |  | speech-in-noise |  | 26 | 28.5 | 740 |

For our dataset (SparrKULee), we conducted an EEG experiment in which 85 participants were recruited and presented with speech stimuli for a duration ranging between 90 and 150 minutes, divided into 6 to 10 recordings (i.e., an uninterrupted period in which a participant listens to a stimulus), totaling 168 hours of EEG data. A general summary can be found in table 2. To validate the obtained dataset, we employed state-of-the-art linear^2,8,18^ and deep learning models^12^, in participant-specific and participant-independent training scenarios. These models can serve as benchmarks for comparison in future research. Our dataset is publicly available on the RDR KU Leuven website.

**Table 2.** Detailed information about the dataset.

| Parameters | Values |
| --- | --- |
| Number of participants | 85 |
| Minutes data per participant | 90 to 150 |
| Number of sessions for each participant | 1 |
| Number of trials per session | 6 to 10 |
| Original sampling rate | 8192 Hz |
| Provided sampling rate | 1024 Hz |
| Number of channels | 64 |

## Methods

We define a *trial* as an uninterrupted recording lasting around 15 minutes. We define a *session* as the complete set of trials and pre-screening activities that a participant underwent from the moment they entered the room until the moment they left. *Stimulus*, in our study, refers to the speech audio files that we presented to the participants during the experiment, which were designed to elicit specific responses from their brains. Figure 1 provides a high-level overview of the different parts of a session.

**Figure 1.**
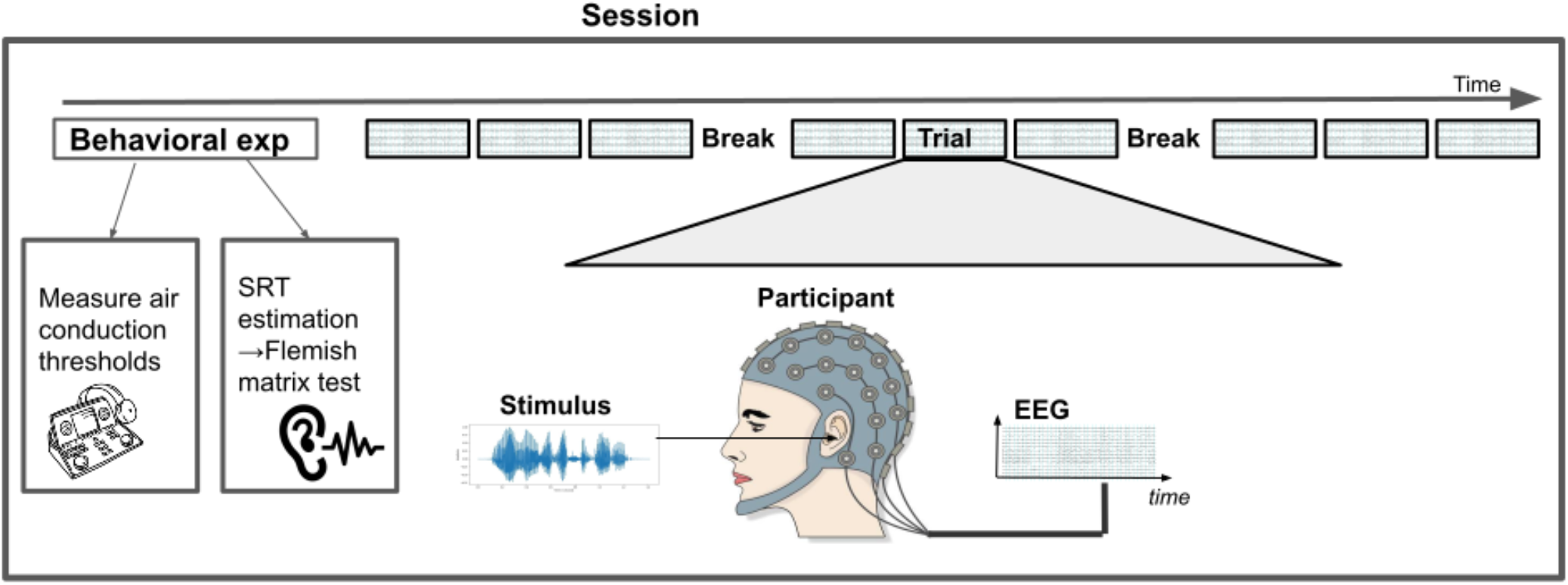
Overview of a session. First, the participant underwent behavioral experiments: air conduction thresholds were measured using the Hughson-Westlake method and the Flemish MATRIX test estimated the Speech reception threshold (SRT). Following the Flemish MATRIX test, the EEG part of the study started, consisting of multiple trials of EEG recording. A trial is defined as an uninterrupted EEG measurement when a stimulus is playing. In this study, trials were approximately 15 minutes in length. After three trials, the participants were offered the option to take a short break.

### Participants

Between October 2018 and September 2022, data were collected from 85 participants (74 female/11 male, 21.4 ±1.9 years (sd)). Inclusion criteria for this study were young (18-30 years), normal-hearing adults (all hearing thresholds <= 30 dB SPL, for 125-8000 Hz), with Dutch/Flemish as their native language. Before commencing the EEG experiments, participants read and signed an informed consent form approved by the Medical Ethics Committee UZ KU Leuven/Research (KU Leuven, Belgium) with reference S57102. All participants in this dataset explicitly consented to share their pseudonymized data in a publicly accessible dataset. This dataset is a subset of our larger proprietary dataset containing data from participants that did not give consent to share their data. Additionally, the participants completed a questionnaire requesting general demographic information (age, sex, education level, handedness^19^) and diagnoses of hearing loss and neurological pathologies. Participants indicating any neurological or hearing-related diagnosis were excluded from the study. Last, the medical history and the presence of learning disabilities were questioned as research has shown that serious concussions, the medication used to treat, for example, insomnia^20^, and learning disabilities such as dyslexia can affect brain responses^21,22^. Therefore this information was used to screen out participants with possibly divergent brain responses.

### Behavioral Experiments

First, we measured the air conduction thresholds using the Hughson-Westlake method^23^ for frequencies from 125 to 8000 Hz (see Figure 2). Participants with hearing thresholds > 30 dB SPL were excluded.

**Figure 2.**
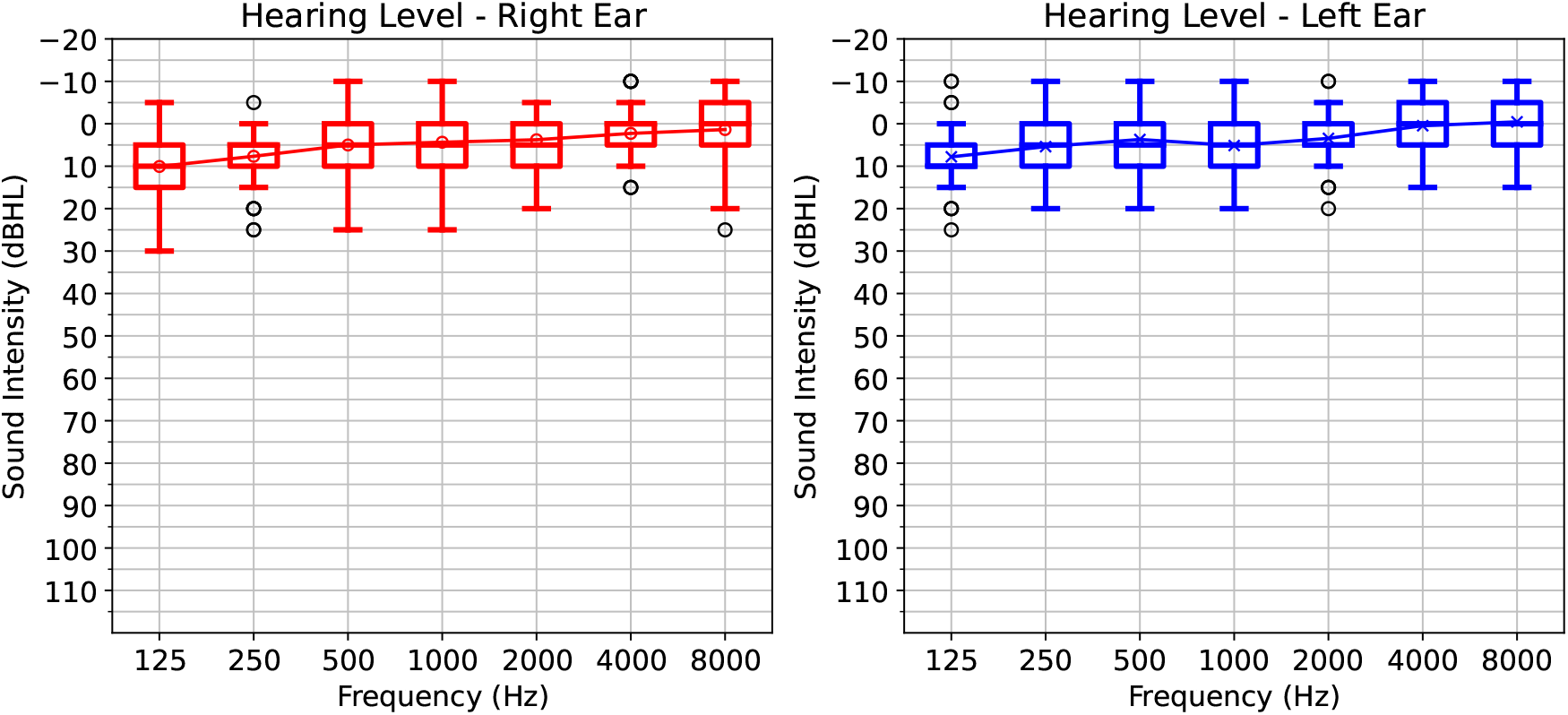
Air conduction thresholds (in dB hearing level (HL)) of the participants.

Secondly, we used the Flemish Matrix test^24^ to determine each participant’s speech reception threshold (SRT, the signal-to-noise ratio (SNR) at which 50 % speech understanding is achieved). The test consisted of 3 lists (2 for training, 1 for evaluation) of 20 sentences following the adaptive procedure of Brand et al.^25^. Each sentence has a fixed syntactic structure of 5 words: name, verb, numeral, color and object [e.g. “Lucas telt vijf gele sokken” (“Lucas counts five yellow socks”)]. After each sentence, participants were asked to indicate the heard sentence using a 5×11 matrix containing ten possibilities for each word and a blank option. The order of the three lists was randomized across participants. The last SNR value was used as an estimate of the SRT. The lists were presented to the participants using electromagnetically shielded Ethymotic ER-3A insert phones, binaurally at 62 dBA for each ear. Luts et al.^24^ present the list to the participants monoaurally to the best ear and obtain an average SRT of −8.7*dBSNR* when using the results of the third list of the adaptive procedure. During the first repetitions, they report a significant training effect, which disappears starting from the third repetition. In our setup, binaural stimulation was chosen to be close to our EEG data acquisition setup. Figure 3 shows the histogram of the obtained SRT over participants in our study. Participants scored an average value of −8.9*dB±* 0.6(*sd*), similar to results obtained by Luts et al.^24^.

**Figure 3.**
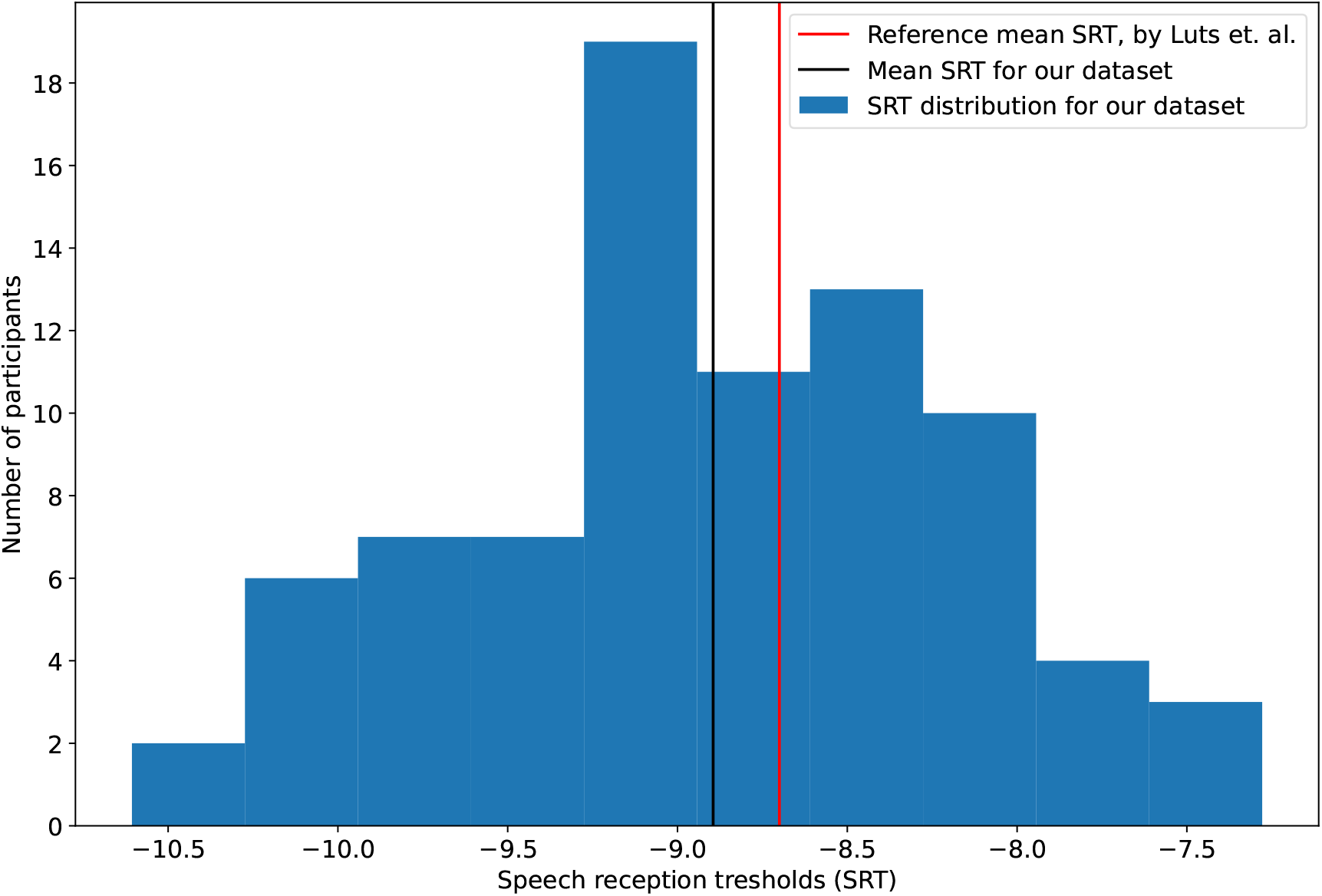
Histogram of the speech reception threshold (SRT), as determined by the matrix test

### EEG data acquisition

All recording sessions were conducted at the research group ExpORL of KU Leuven, in a triple-walled, soundproof booth equipped with a Faraday cage to reduce external electromagnetic interference. Participants were instructed to listen to the speech while seated and minimize muscle movements. They were seated in a comfortable chair in the middle of the booth.

We recorded EEG using a BioSemi ActiveTwo system with 64 active Ag-AgCl electrodes and two additional electrodes for the common electrode (CMS) and current return path (DRL). In addition, two mastoid electrodes and the BioSemi head caps were used, containing electrode holders placed according to the 10-20 electrode system.

To ensure proper electrode placement for each participant, we first measured their head size (from nasion to inion to nasion) and selected an appropriate cap. Mastoid locations were scrubbed with Nuprep and cleaned with alcohol gel. The mastoid electrodes were then attached using stickers and held with tape.

The electrode cap was placed on the participant’s head from back to front, with ears placed through gaps in the cap. The closing tape at the bottom was secured, and a visual assessment was performed to ensure proper fit. The cap was adjusted so that the distance between the nasion and the electrode Cz, the inion and the electrode Cz were equal, and the distance between the left and right ears and the Cz electrode. Electrode gel was applied to the cap holes, and the electrodes were placed gently. The battery, electrode cables and mastoid electrodes were attached to the BioSemi AD-box. The participant was then instructed to sit still while EEG was recorded. The subjects were told to keep their eyes open during the measurement. If necessary, the additional gel was applied to poorly behaving electrodes, and the electrode offset was checked to ensure proper connection. All offsets were ideally between +20 and -20 mV.

The EEG recordings were digitized at a sampling rate of 8192 Hz and stored on a hard disk using the BioSemi ActiView software.

### EEG experiment

All participants listened to 6, 7, 8 or 10 trials, each of approximately 15 minutes. The order of all the trials was randomized per participant. After each trial, a question about the stimulus content was asked to determine attention to and comprehension of the story. As the questions were not calibrated, they merely motivated the participant to pay attention to the stimulus. After three trials, the participants were asked if they wanted to have a short break. Table 3 shows an overview of the experiment and timing.

**Table 3.**
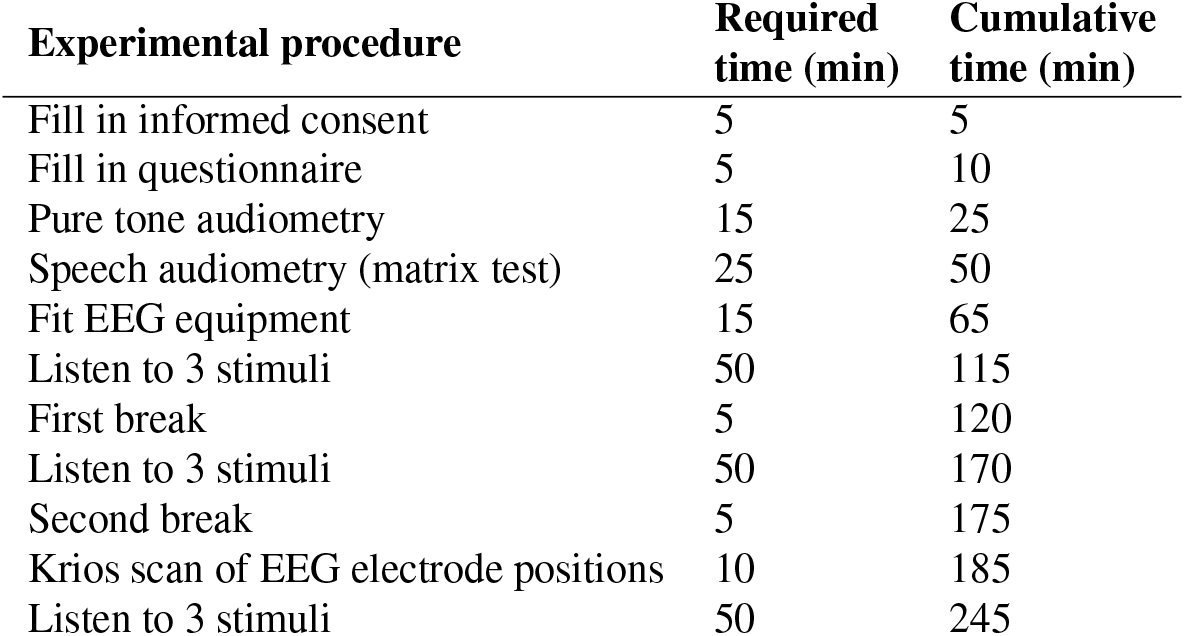
Overview of the experimental procedure.

We used different categories of stimuli:

- **Reference audiobook** to which all participants listened, made for children and narrated by a male speaker. The length of the audiobook is around 15 minutes.
- **Audiobooks** made for children or adults. To keep the trial length around 15 minutes, some audiobooks were split into different *parts* when the length exceeded 15 minutes.
- **Audiobooks with noise** made for children to which speech-weighted noise was added, as explained below, to obtain an SNR of 5 *dB*.
- **Podcasts** from the series’ Universiteit van Vlaanderen’ (University of Flanders)^26^. Each episode of this podcast answers a scientific question, lasts around 15 minutes, and is narrated by a single speaker.
- **Podcasts with video** from the series’ Universiteit van Vlaanderen’ (University of Flanders)^26^, while video material of the speaker was shown. The video material can be found on the website of Universiteit van Vlaanderen for each podcast separately.

The Podcasts and Podcasts with video were dynamically range compressed by the producers of the stimuli, while the audiobooks were not.

The dataset collection consists of two main session types: ses-shortstories01 and ses-varyingstories, differing in the presented stimuli. Each participant undertook one session. An overview of the experiment and timing can be found in table 3, while figure 4 summarizes which stimuli were used for each participant in each session.

**Figure 4.**
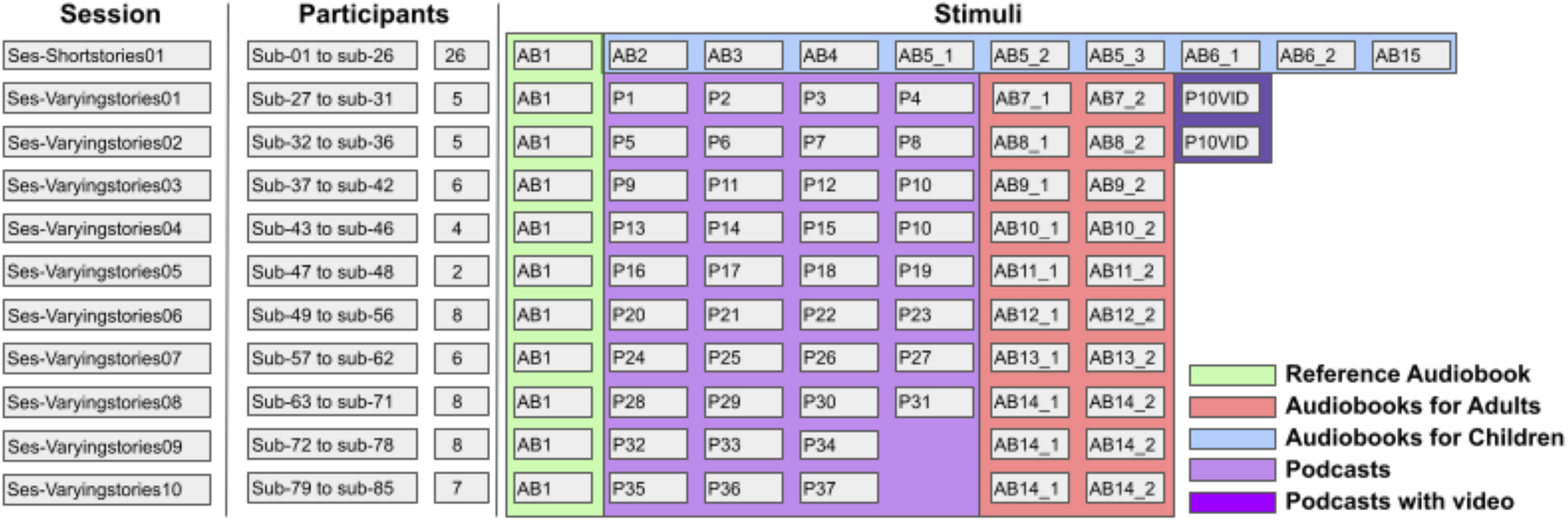
Overview of all the stimuli that were presented, per participant. AB=audiobook, P=podcast. Audiobooks and podcast are numbered. The subscript *_1/2/3* indicate different parts of the same audiobook, each around 15 minutes in length.

#### Ses-shortstories01

For this session type, data from 26 participants is available. It includes ten different parts of audiobooks for children. Two audiobooks, audiobook_1 and audiobook_4, are narrated by male speakers, the other by female speakers. audiobook_3, audiobook_5 and audiobook_6 are narrated by the same speaker. Two out of ten trials were randomly chosen for each participant and presented in speech-weighted-noise (SNR = 5dB). Additionally, 3 subjects listened to a different version of audiobook_1. For this experiment, the audiobook was cut in 2 halves (audiobook_1_1, audiobook_1_2 respectively), and a pitch-shifted version was used created for each half (audiobook_1_1_shifted, audiobook_1_2_shifted, respectively). More information about the pitch shifting and additional experiments can be found in the work of Algoet et al.^27^. Finally, there was one control condition in which the first 5 minutes of audiobook_1 were presented to a subject who had no insertphones inserted (audiobook_1_artefact]).

#### Ses-varyingstories

For the ses-varyingstories type, data from 59 participants are available. Ses-varyingstories had a fixed reference audiobook_1 (which was presented to all subjects), an audiobook of around 30 minutes split into two parts, and three to five different podcasts per participant, chosen to keep an even distribution of the sex of the speaker. The stimuli were changed every 2 to 8 participants.

### Stimulus preparation

All stimuli were stored at a sampling rate of 48*kHz*. For each stimulus file, a trigger file was generated. These triggers were sent from the stimulation equipment (RME soundcard) to the BioSemi. Triggers were generated every second in the form of a block wave. At every second and the beginning and end of the recording, a block pulse with a width of 1 ms is inserted. Based on the stimulus, speech-shaped noise was created at the same root-mean-square value (RMS) as the stimulus. The noise was created by taking white noise and changing the spectrum of the white noise to the spectrum of the speech, and then matching the RMS value of the original stimulus file.

Afterward, using one noise file for each RMS value, the stimuli were calibrated with a type 2260 sound-level pressure meter, a type 4189 0.5-in. microphone, and a 2-cm3 coupler (Bruel & Kjaer, Copenhagen, Denmark).

The auditory stimuli were presented using a laptop connected to an RME Hammerfall DSP Muliface II or RME Fireface UC soundcard, using the APEX software platform^28^ and electromagnetically shielded Ethymotic ER-3A insert phones, binaurally at 62 dBA for each ear.

### Krios data

We acquired a 3D-scan of the configuration of the EEG caps for all participants, using a Polaris Krios scanner (NDI, Canada), which scans all the electrodes, using a probe to mark three reference points: at the nasion and the height of the tragus at both sides. The Polaris Krios scanner is based on optical measurement technology and uses light reflected by markers to determine the position coordinates.

### EEG data preprocessing

Besides the raw EEG recordings, we also provide EEG with commonly used preprocessing steps applied. All steps were conducted in Python 3.6, and the code for preprocessing is available on our GitHub repository (https://github.com/exporl/auditory-eeg-dataset). First, EEG data was high-pass filtered, using a 1st-order Butterworth filter with a cut-off frequency of 0.5 Hz. Zero-phase filtering was conducted by filtering the data forward and backward. Subsequently, the EEG was downsampled from 8192 Hz to 1024 Hz and eyeblink artifact removal was applied to the EEG, using a multichannel Wiener filter^30^. Afterward, the EEG was re-referenced to a common average, and finally, the EEG was downsampled to 64 Hz.

### Speech stimuli preprocessing

The initial sampling frequency of the stimuli was 48kHz. We provide a script to calculate the envelope using a gammatone filterbank^31^ with 28 subbands. Each subband envelope was calculated by taking the absolute value of each sample, raised to the power of 0.6. A single envelope was obtained by averaging all these subbands^32^. Then, the envelope was downsampled to 64 Hz.

## Data Records

All data were organized according to EEG-BIDS^33^, an extension to the Brain Imaging Directory Structure (BIDS)^34^ for EEG data. EEG-BIDS allows storing EEG data with relevant extra information files, e.g., about the experiment, the stimuli and the triggers, enabling quick usage of the data and linking the auditory stimuli to the raw EEG files. A schematic overview of our repository is shown in Figure 5. The dataset consists of 3 parts: (1) raw data, in a folder per participant, (2) the auditory stimuli, in zipped Numpy (.*npz*)^35^ format (3) the preprocessed data records, as described above, in the *derivatives* folder.

**Figure 5.**
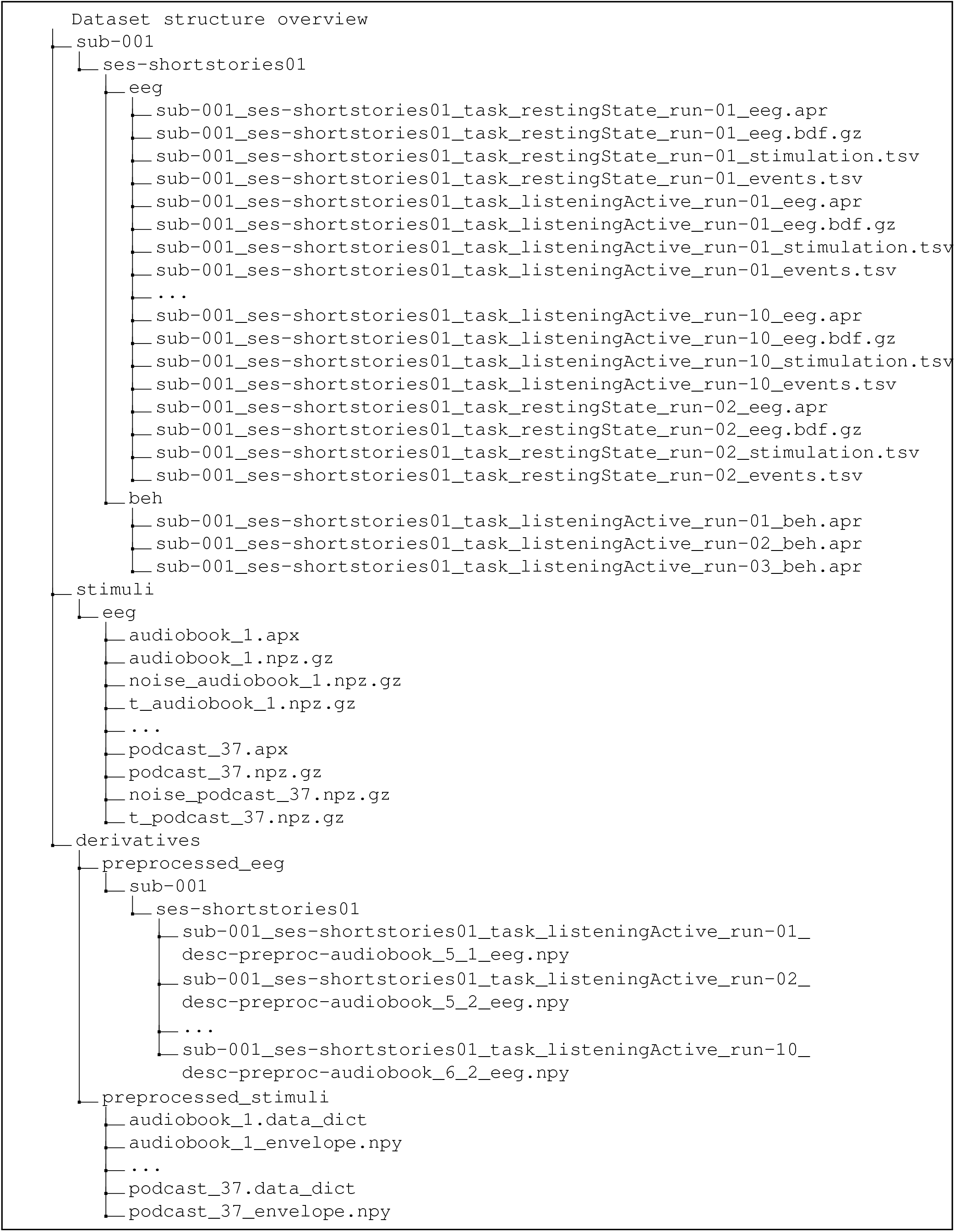
Tree depicting the structure of our dataset. All data have been structured according to the EEG-BIDS standard.

### Raw data

The raw data was structured in a folder per participant. For each participant (1 to 85), a folder `sub-xxx` is available in the root folder. In this folder, there is a folder indicating the session, which can be either `ses-shortstories01` or `ses-varyingstoriesxx` (*xx* = 01 … 09).

Each session folder contains a subfolder `beh`, containing the results of the behavioral matrix SRT estimation. These files were named according to the participant, the session, the task, which is always *listeningActive*, and the behavioral experiment run, which goes from 1 to 3.

The data of the EEG experiment was stored as a subfolder in the session folder, named `eeg`. The EEG experiment data were named according to the participant, the session, the task and the run. When the participant listened to a stimulus, the task was `listeningActive`. When the participant listened to silence, which happened at the start and end of the experiment, the task was`restingState`. The run suffix chronologically numbers the different trials starting at 01. Each trial has four corresponding files, differing only in their ending, after the run suffix: (1) raw gzipped file of EEG data in BioSemi Data Format (BDF), sampled at 8192 Hz, ending in `eeg.bdf.gz`, (2) a descriptive apr file `eeg.apr`, containing extra information about the experiment, such as the answers to the questions that were asked, (3) stimulation file to link EEG to the corresponding stimulus `stimulation.tsv` and (4) `events.tsv`, which describes which stimuli were presented to the participants at which time.

### Stimuli

All the stimuli are saved in the folder `stimuli/eeg`. For each stimulus, we provide four corresponding files, stored in the *npz* format with additional gzipping to reduce storage, which is easily readable in Python: (1) the stimulus, stored at 48 kHz `stimulusName.npz.gz`, (2) the associated noise file `noise_stimulusName.npz.gz`, (3) the associated trigger file `t_stimulueName.npz.gz` and (4) the experiment description file `stimulusName.apx`.

The stimuli were named according to their type: either `audiobook_xx` or `podcast_xx`, where *xx* indexes unique stimuli. Whenever an audiobook was split into multiple consecutive parts, an extra suffix denotes which part of the audiobook is referred to.

### Preprocessed data

For all data, we also provide a preprocessed, downsampled version of the data. These data can be found in the `derivatives/preprocess` folder. Similar to the raw data, the preprocessed data was structured in a folder per participant, per session, which could be either ses-shortstoriesxx or ses-varyingstoriesxx. The preprocessed files derive their name from the raw EEG file used to create the preprocessed version. To avoid confusion, a suffix `desc-preproc` was added, such that no two files have the same name. After the `desc-preproc` suffix, the name of the stimulus the participant listened to was added to facilitate linking the EEG brain response to the auditory stimulus for downstream tasks.

## Technical Validation

In order to demonstrate the validity of the data, we conducted several experiments on the preprocessed version of the proposed dataset. The code to obtain these results can be found online: https://github.com/exporl/auditory-eeg-dataset.

### Additional preprocessing

For all our experiments, we split each trial into a training, validation and test set, containing respectively 80%, 10% and 10% of each trial for each participant. The train, validate and test set do not overlap, so the test set remains unseen for all the models.

Before usage, we normalized each trial by computing the mean and standard deviation for each of the 64 EEG channels and the envelope stimulus on the training set. We then normalized the train, validation and test set by subtracting from each trial the mean and dividing by the standard deviation computed on the train set.

### Linear forward/backward modeling

To show the validity of the data, we trained participant-specific linear forward and backward models^1,2^ (i.e., models that predict EEG from the stimulus envelope and the stimulus envelope from the EEG, respectively). The backward model was used to detect neural tracking in each recording, i.e., that the speech envelope can effectively be decoded for each participant/story compared to a null distribution of random predictions. The forward model was used to visualize the EEG channels for which the stimulus-related activity can be best predicted.

#### Model training

The models were trained based on the recommendations of Crosse et al. (2021)^18^. The backward model weights were obtained similarly by equation 1:

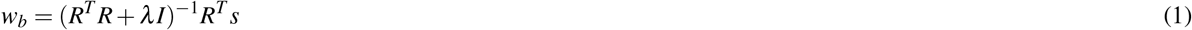

Where *R* is a matrix consisting of time-lagged versions of the EEG, *s* is the stimulus envelope and *λ* is the ridge regression parameter.In a similar fashion, equation 2 was used to obtain the forward model weights:

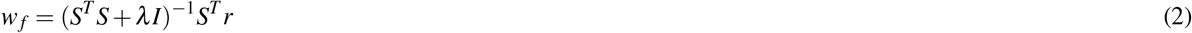

Where *S* is a matrix consisting of time-lagged versions of the stimulus envelope, *r* is a matrix containing the EEG response, and *λ* is the ridge regression parameter.

Both models had an integration window from -100ms to 400ms. Following the recommendations of Crosse et al. (2021)^18^, leave-one-out cross-validation was performed on the recordings in the training set to determine the optimal ridge regression parameter (*λ*) from a list of values (10^*x*^ for *x* = [− 6, − 4, − 2, 0, 2, 4, 6]). Correlations scores were averaged across folds and channels, after which the *λ* is chosen, corresponding to the highest correlation value.

To evaluate the performance of both models, the Pearson correlation between the predicted and true data was calculated on the test set. In order to detect neural tracking, we followed the procedure of Crosse et al. (2021)^18^. For each recording in the test-set, the predictions are (circularly) shifted in time by a random amount *N* = 100 times. By correlating these shifted predictions to the actual signal, a null distribution was constructed for each participant. The 95th percentile of this null distribution was compared to the mean of the obtained scores on the test sets.

The analysis of EEG neural responses is typically performed in specific filter bands. For auditory EEG, the research typically focuses on the Delta band (0.5 − 4 *Hz*) and the Theta band (4 − 8 *Hz*)^2,36–38^. We investigated the effect of filtering the EEG and envelope in different bands: Delta (0.5 − 4*Hz*), Theta (4 − 8*Hz*), Alpha (8 − 14*Hz*), Beta (14 − 30*Hz*) and Broadband (0.5 − 32*Hz*). A 1st order Butterworth filter was chosen for each of the proposed filtering bands.

The model training and evaluation were performed in Python using Numpy^35^ and Scipy.

#### Analysis

Using the linear backward model, we were able to detect neural tracking for all participants. In 11 of the 666 recordings, we were not able to detect neural tracking in any frequency band with the linear decoder. These recordings are listed in table 4. The results per frequency band are shown in Figure 6. As previously shown by Vanthornhout et al.^2^, the optimal performance was reached when filtering in the delta-band (0.5 − 4 *Hz*). While correlations are hard to compare between studies because they are heavily influenced by the measurement paradigm, subject selection, preprocessing and modeling choices, the correlations we found for the delta band are roughly in line with previous studies (median correlation between 0.1-0.2^1,2^).

**Table 4.** Recordings where no significant tracking was found with the linear backward model.

| Subject | Stimulus |
| --- | --- |
| sub-002 | audiobook_1_artefact |
| sub-011 | audiobook_6_1 |
| sub-051 | audiobook_12_1 |
| sub-051 | audiobook_12_2 |
| sub-051 | podcast_23 |
| sub-054 | audiobook_12_2 |
| sub-056 | podcast_22 |
| sub-060 | podcast_24 |
| sub-064 | audiobook_14_2 |
| sub-064 | podcast_30 |
| sub-076 | audiobook_14_1 |

**Figure 6.**
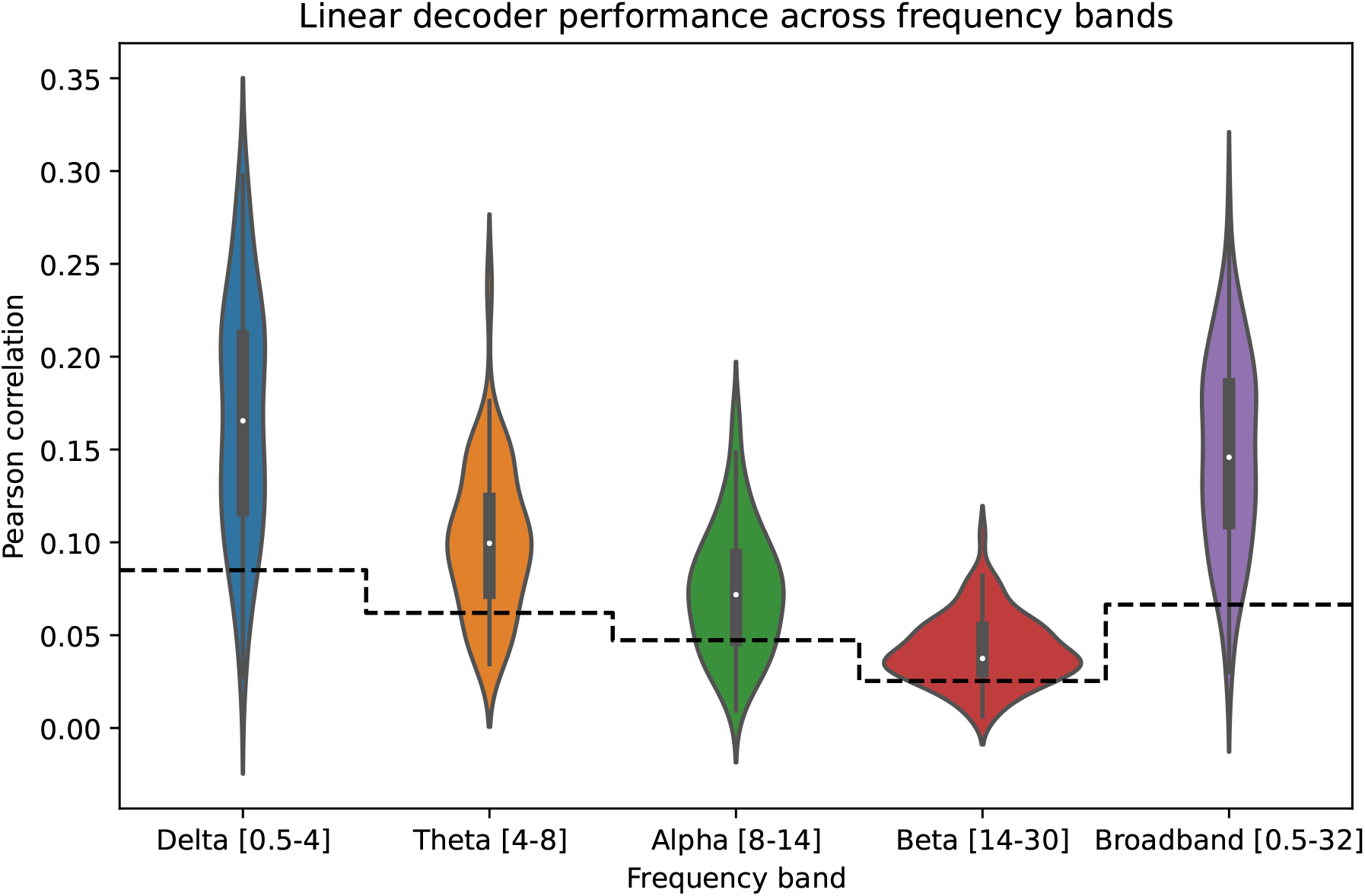
Results of the linear backward model for different frequency bands. Each point in the boxplot is the correlation between the predicted speech envelope and stimulus envelope for one participant, averaged over recordings. Separate models were trained for each participant and frequency band (Delta (0.5 − 4*Hz*), Theta (4 − 8*Hz*), Alpha (8 − 14*Hz*), Beta (14 − 30*Hz*) and Broadband (0.5 − 32*Hz*)). Highest correlations were obtained in the delta band and decreased when going to higher frequency bands. The dashed line represents the significance level (*α*=0.05)

We compared the linear backward model performance across all stimuli and stimuli types (audiobooks vs. podcast, excluding the audiobook_1 shifted and artifact versions) in the delta-band. The results are visualized in Figure 7 and Figure 8, respectively. Note that there is a large variability in decoding scores within and between stimuli. Additionally, a significant difference was found between the audiobook and podcast stimuli (0.184 vs. 0.133 median Pearson correlation, MannWhitneyU test: *p <* 10^−9^).

**Figure 7.**
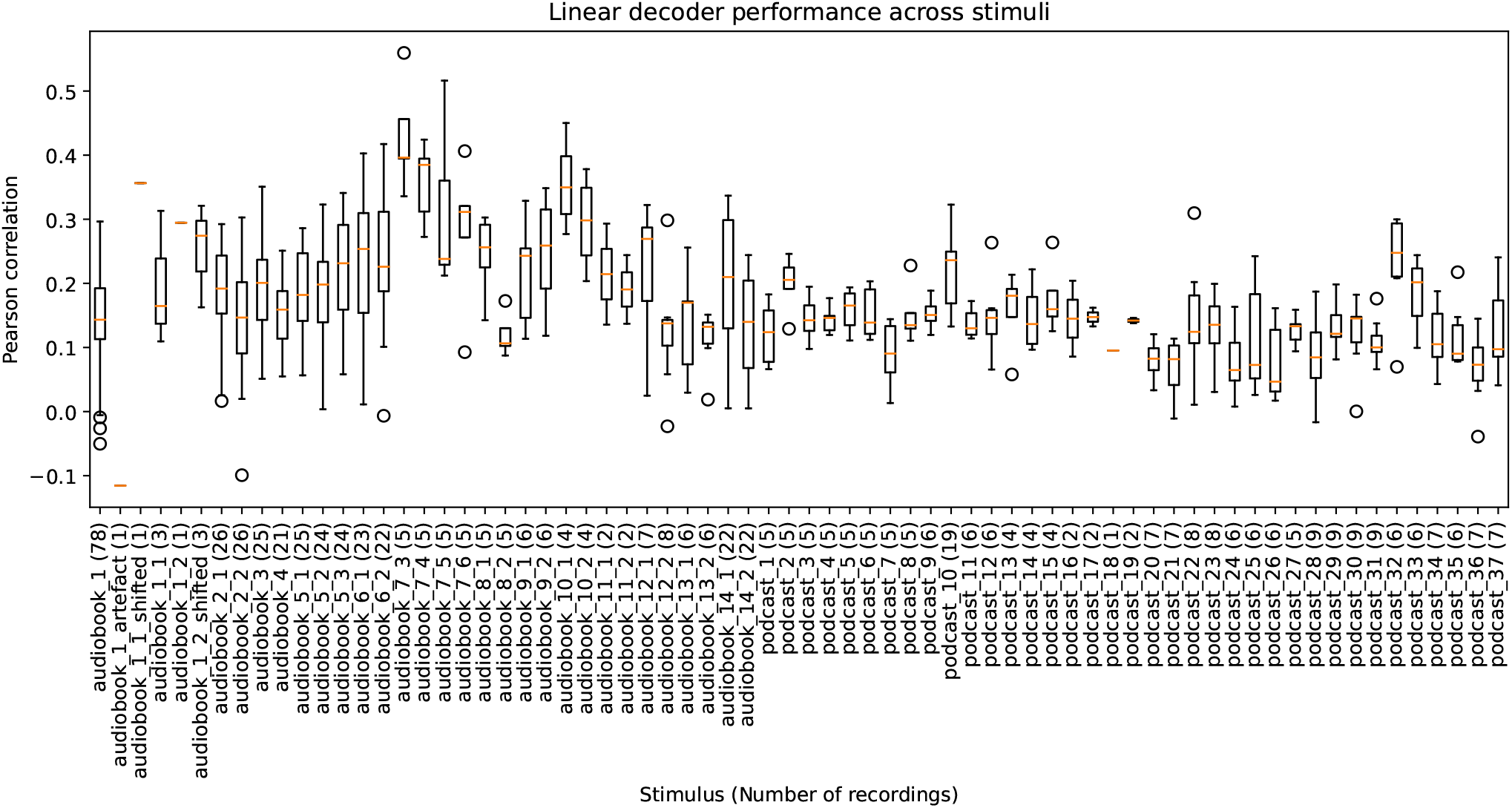
Results of the linear backward model for the different stimuli in the dataset. One model is trained per participant. Each point in the boxplot is the correlation between the predicted speech envelope and stimulus envelope for one recording. Data was filtered in the delta band (0.5 − 4*Hz*). There is high variability across participants and stimuli.

**Figure 8.**
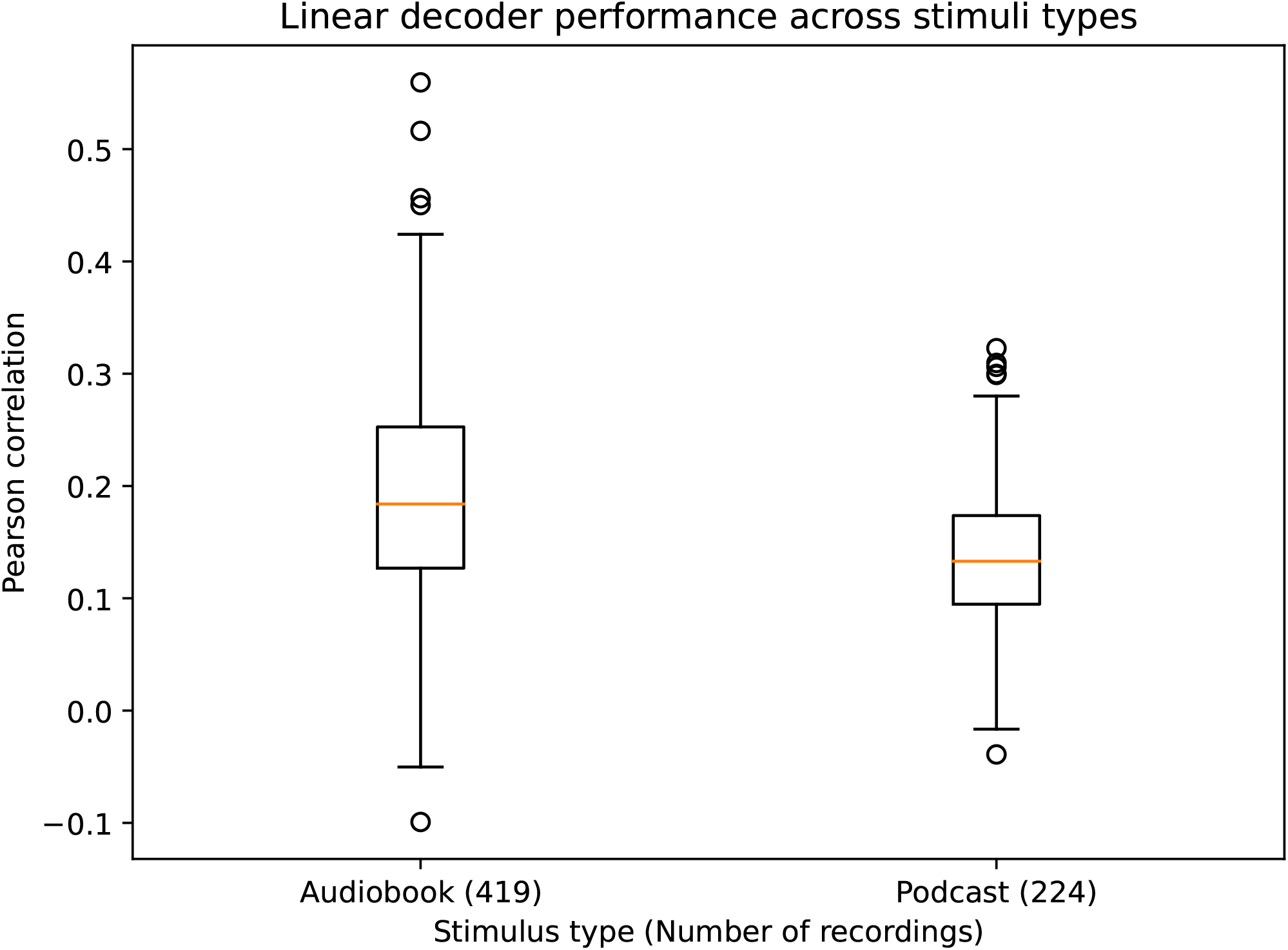
Results of the linear backward model for the different stimuli in the dataset. One model is trained per participant. Each point in the boxplot is the correlation between the predicted speech envelope and stimulus envelope for one participant, averaged across recordings. Significantly higher correlations were obtained for the audiobooks (0.184 vs. 0.133 median Pearson correlation, MannWhitneyU test: *p <* 10^−9^).

For the forward model, we show topomaps averaged across participants for each frequency band and stimulus type in Figure 9. As with the backward model, we observed the highest correlations between predicted and actual EEG signals in the delta band. The highest correlations were obtained for the channels in the temporal and occipital regions.

**Figure 9.**
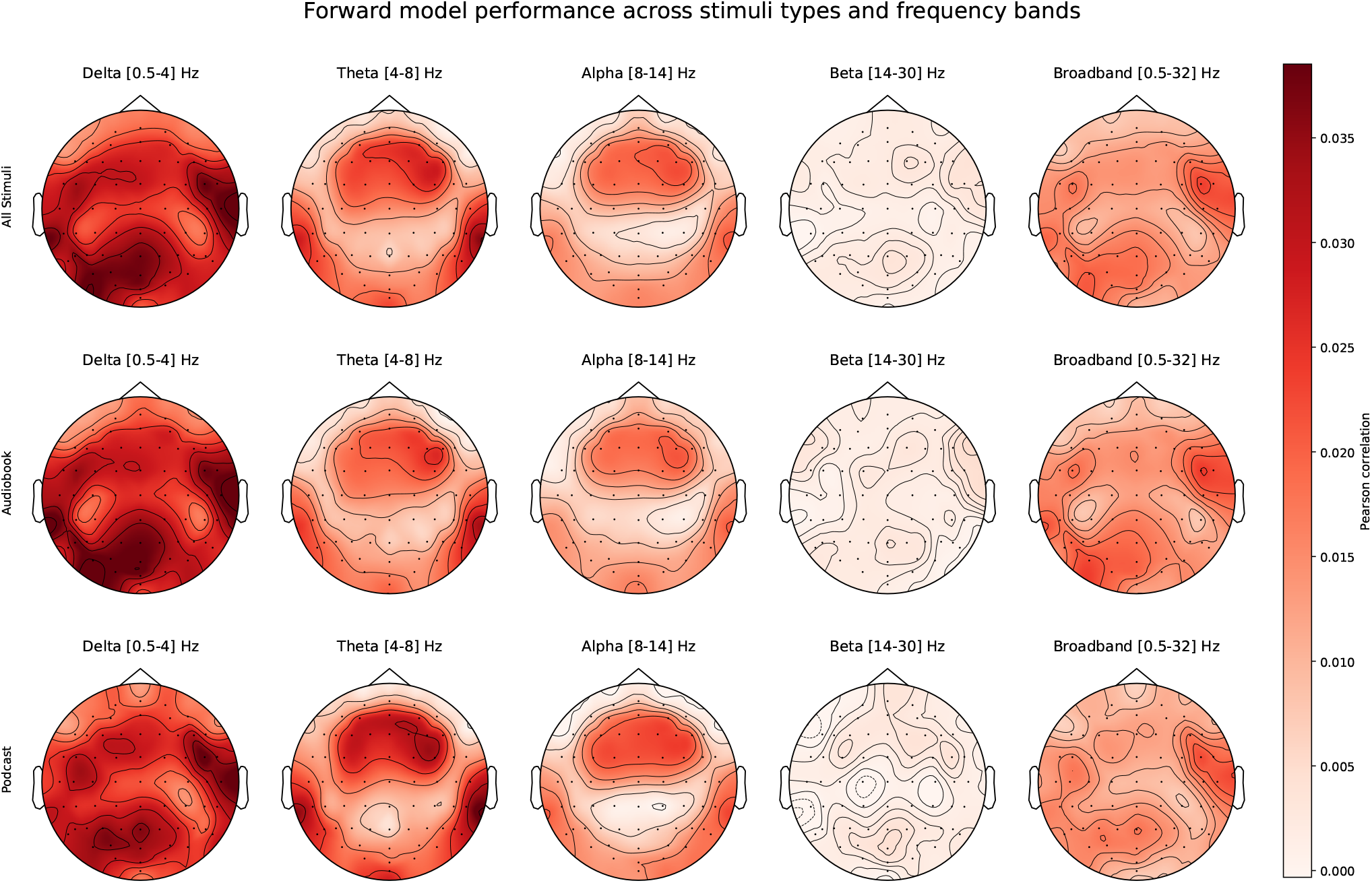
Results of the forward linear model for different stimuli types and frequency bands. For each channel, the correlation between actual and predicted EEG is shown and averaged across participants. One model is trained per participant. The highest correlations are obtained in the delta-band for the channels in the temporal and occipital region.

### Non-linear models - Match-mismatch paradigm

For the non-linear models, we used the match-mismatch paradigm^7,12^. In this paradigm, the models are given three inputs: a segment of the EEG recording, the time-matched stimulus envelope segment, and a mismatched (imposter) stimulus envelope segment. As specified by^10^, the imposter was taken 1s after the matched stimulus envelope segment. If extracting an imposter (at the end of each set) was impossible, the segment was discarded from the dataset. We extracted overlapping windows with 80% overlap. We included an analysis using a dilated convolutional model^12^ to show typical match-mismatch performance across different input segment lengths.

#### Model training

The dilated convolutional network consists of four steps. First, the EEG channels are combined, from 64 to 8, using a 1D convolutional layer with a kernel size of 1 and a filter size of 8. Second, there are *N* dilated convolutional layers with a kernel size of *K* and 16 filters. These *N* convolutional layers are applied to both EEG and envelope stimulus segments. After each convolutional layer, a rectified linear unit (ReLU) is applied. Both stimulus envelope segments share the weights for the convolutional layers. After these non-linear transformations, the EEG is compared to both stimulus envelopes, using cosine similarity. Finally, the similarity scores are fed to a single neuron, with sigmoid non-linearity, to create a prediction of the matching stimulus segment.

The model was implemented in Tensorflow and used the Adam optimizer, with a learning rate of 0.001 and binary-cross entropy as the loss function. Models were trained for a maximum of 50 epochs, using early stopping based on the validation loss, with a patience factor of 5. We trained the models with an input segment length of 5 seconds and in a participant-independent way, i.e., all participant data was given simultaneously to the model. We report results for input testing lengths 1, 2, 3, 5, and 10 s. Since the trained dilation model does not have fixed input lengths, we used the same model with different input lengths.

#### Analysis

The results of this analysis can be seen in figure 10. The accuracy of the model increased with longer window lengths. We see the same trend as in^12^. In order to test the generalizability of the model, we also tested the model with an arbitrarily chosen mismatch segment, as opposed to the fixed 1 second. There was no significant difference between these two testing conditions, which is in line with the experiment as conducted in^39^.

**Figure 10.**
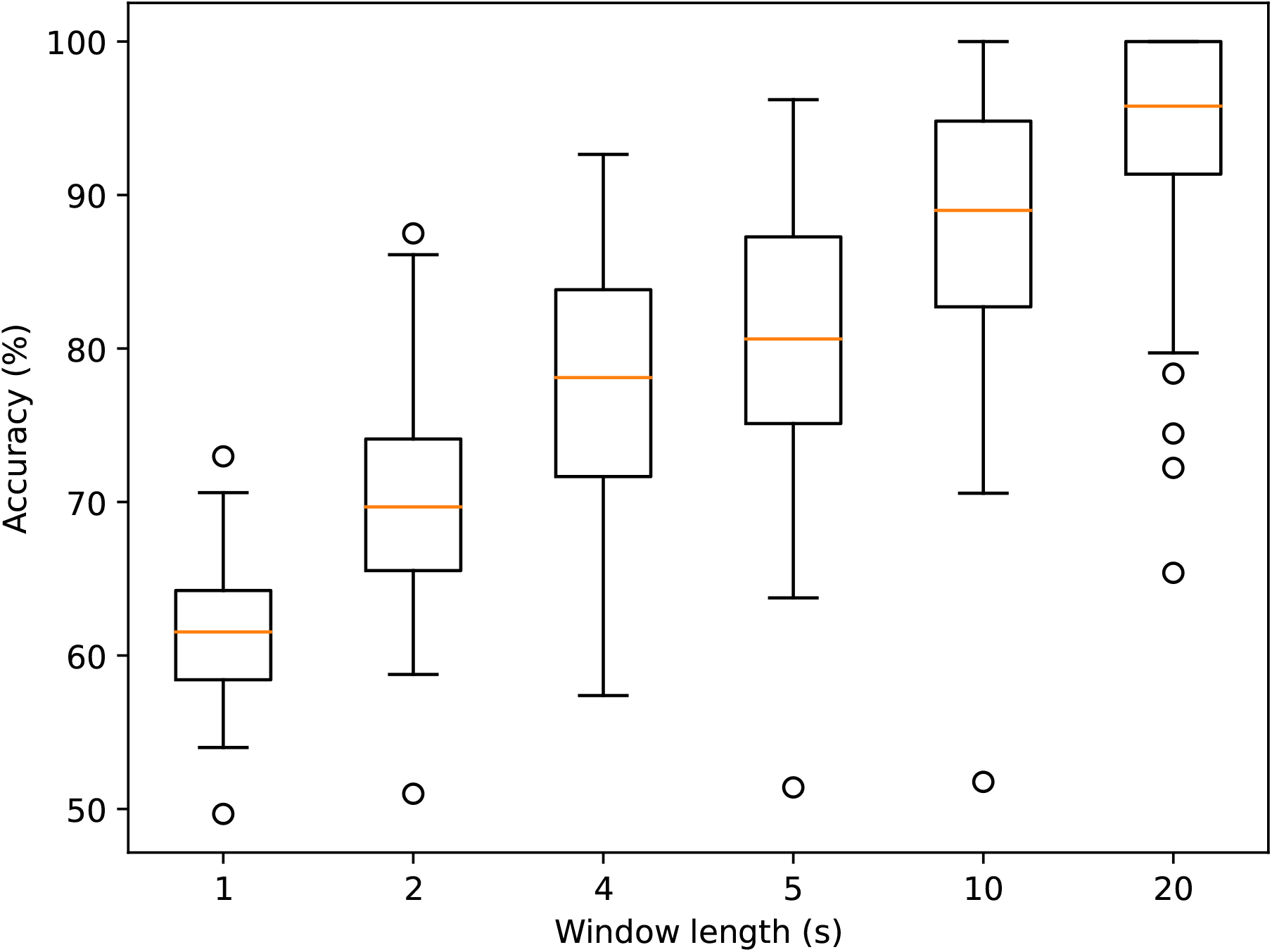
Results of the non-linear dilation model, in the match-mismatch paradigm. Each point in the boxplot is the match-mismatch accuracy for one participant, averaged across recordings. The imposter envelope segment starts one second after the end of the true segment. One model was trained across all participants.

## Usage Notes

The stimuli included in the dataset were saved in Numpy array format^35^. *AB*_1_, *AB*_3_, *AB*_*xp*1_ and *AB*_*xp*2_ for *x* = 7…14 originate from the Radioboeken project of deBuren (https://soundcloud.com/deburen-eu/). Podcasts were obtained from Universiteit van Vlaanderen (https://www.universiteitvanvlaanderen.be). All stimuli in the dataset can only be used/shared for non-commercial purposes. When republishing (adaptations of) the stimuli, explicit permission should be acquired from the original publishing organization(s) (i.e., deBuren or Universiteit van Vlaanderen).

The dataset is available on the RDR KU Leuven platform https://rdr.kuleuven.be/dataset.xhtml?persistentId=doi:10.48804/K3VSND under an Attribution-NonCommercial 4.0 International License (CC BY-NC 4.0). Due to privacy concerns, access to part of the data is restricted. Readers requesting access should mail the corresponding authors, stating what they want to use the data for. Access will be granted to non-commercial users, complying with the CC-BY-NC-4.0 license.

## Code availability

All code used for the technical validation can be found online: https://github.com/exporl/auditory-eeg-dataset. We used the mne-python library^40^.

For using the data, we recommend using the code on our GitHub repository to get started, which consists of two main parts: (1) code to create the preprocessed eeg and preprocessed stimuli from the raw data and (2) code to perform the experiments as discussed in the technical validation. The README file contains detailed technical instructions.

## Acknowledgements

The research conducted in this paper is funded by KU Leuven Special Research Fund C24/18/099 (C2 project to Tom Francart and Hugo Van hamme), by two PhD grants (1S89622N,1SB1421N) of the Research Foundation Flanders (FWO) and from the European Research Council (ERC) under the European Union’s Horizon 2020 research and innovation program (grant agreement No 637424, ERC Starting Grant to Tom Francart).

The authors thank Amelie Algoet, Jolien Smeulders, Lore Kerkhofs, Sara Peeters, Merel Dillen, Ilham Gamgami, Amber Verhoeven, Vitor Vasconselos, Jard Hendrickx, Lore Verbeke and Ana Carbajal Chavez for their help with data collection.

## Author contributions statement

B.A., W.V., L.B., H.V.h and T.F conceived the experiments, B.A., L.B., M.G. and W.V. conducted/supervised the experiments, B.A. and L.B analysed the results. All authors reviewed the manuscript.

## References

1. Crosse, M. J., Di Liberto, G. M., Bednar, A. & Lalor, E. C. The multivariate temporal response function (mTRF) toolbox: A MATLAB toolbox for relating neural signals to continuous stimuli. Front. Hum. Neurosci. 10, 1–14, 10.3389/fnhum.2016.00604 (2016).

2. Vanthornhout, J., Decruy, L., Wouters, J., Simon, J. Z. & Francart, T. Speech Intelligibility Predicted from Neural Entrainment of the Speech Envelope. JARO - J. Assoc. for Res. Otolaryngol. 19, 181–191, 10.1007/s10162-018-0654-z (2018).

3. Iotzov, I. & Parra, L. C. EEG can predict speech intelligibility. J. Neural Eng. 16, 036008, 10.1088/1741-2552/ab07fe (2019).

4. Thornton, M., Mandic, D. & Reichenbach, T. Robust decoding of the speech envelope from eeg recordings through deep neural networks. J. Neural Eng. 19, 046007 (2022).

5. Accou, B., Vanthornhout, J., Hamme, H. V. & Francart, T. Decoding of the speech envelope from EEG using the VLAAI deep neural network, 10.1101/2022.09.28.509945 (2022). Pages: 2022.09.28.509945 Section: New Results.

6. Lesenfants, D., Vanthornhout, J., Verschueren, E. & Francart, T. Data-driven spatial filtering for improved measurement of cortical tracking of multiple representations of speech. bioRxiv 10.1101/551218 (2019).

7. de Cheveigné, A. et al. Multiway canonical correlation analysis of brain data. NeuroImage 186, 728–740, https://doi.org/10.1016/j.neuroimage.2018.11.026 (2019).

8. de Cheveigné, A., Slaney, M., Fuglsang, S. A. & Hjortkjaer, J. Auditory stimulus-response modeling with a match-mismatch task. J. Neural Eng. 18, 046040, 10.1088/1741-2552/abf771 (2021).

9. de Taillez, T., Kollmeier, B. & Meyer, B. Machine learning for decoding listeners’ attention from eeg evoked by continuous speech. Eur. J. Neurosci. 51, p10.1111/ejn.13790 (2017).

10. Monesi, M. J., Accou, B., Montoya-Martinez, J., Francart, T. & Hamme, H. V. An LSTM Based Architecture to Relate Speech Stimulus to Eeg. ICASSP, IEEE Int. Conf. on Acoust. Speech Signal Process. - Proc. 2020-May, 941–945, 10.1109/ICASSP40776.2020.9054000 (2020).

11. Monesi, M. J., Accou, B., Francart, T. & Van Hamme, H. Extracting different levels of speech information from eeg using an lstm-based model, 10.48550/ARXIV.2106.09622 (2021).

12. Accou, B., Jalilpour-Monesi, M., hamme, H. V. & Francart, T. Predicting speech intelligibility from eeg using a dilated convolutional network. ArXiv abs/2105.06844 (2021).

13. Ding, N. & Simon, J. Z. Emergence of neural encoding of auditory objects while listening to competing speakers. Proc. Natl. Acad. Sci. 109, 11854–11859, 10.1073/pnas.1205381109 (2012). https://www.pnas.org/doi/pdf/10.1073/pnas.1205381109.

14. Broderick, M. P., Anderson, A. J., Di Liberto, G. M., Crosse, M. J. & Lalor, E. C. Electrophysiological correlates of semantic dissimilarity reflect the comprehension of natural, narrative speech. Curr. Biol. 28, 803–809 (2018).

15. Fuglsang, S. A., Wong, D. D. & Hjortkjær, J. Eeg and audio dataset for auditory attention decoding, 10.5281/zenodo.1199011 (2018).

16. Etard, O. & Reichenbach, T. EEG Dataset for ‘Decoding of selective attention to continuous speech from the human auditory brainstem response’ and ‘Neural Speech Tracking in the Theta and in the Delta Frequency Band Differentially Encode Clarity and Comprehension of Speech in Noise’., 10.5281/zenodo.7086209 (2022).

17. Weissbart, H., Kandylaki, K. & Reichenbach, T. EEG Dataset for ‘Cortical Tracking of Surprisal during Continuous Speech Comprehension’, 10.5281/zenodo.7086168 (2022).

18. Crosse, M. et al. Linear modeling of neurophysiological responses to speech and other continuous stimuli: Methodological considerations for applied research. Front. Neurosci. 15, 10.3389/fnins.2021.705621 (2021).

19. Coren, S. The lateral preference inventory for measurement of handedness, footedness, eyedness, and earedness - norms for young-adults. BULLETIN OF THE PSYCHONOMIC SOCIETY 31, 1–3, 10.3758/bf03334122 (1993).

20. van Lier, H., Drinkenburg, W. H., van Eeten, Y. J. & Coenen, A. M. Effects of diazepam and zolpidem on eeg beta frequencies are behavior-specific in rats. Neuropharmacol. 47, 163–174, https://doi.org/10.1016/j.neuropharm.2004.03.017 (2004).

21. De Vos, A., Vanvooren, S., Vanderauwera, J., Ghesquière, P. & Wouters, J. Atypical neural synchronization to speech envelope modulations in dyslexia. Brain Lang. 164, 106–117, https://doi.org/10.1016/j.bandl.2016.10.002 (2017).

22. Power, A. J., Colling, L. J., Mead, N., Barnes, L. & Goswami, U. Neural encoding of the speech envelope by children with developmental dyslexia. Brain Lang. 160, 1–10, https://doi.org/10.1016/j.bandl.2016.06.006 (2016).

23. Hughson, W., Westlake, H. et al. Manual for program outline for rehabilitation of aural casualties both military and civilian. Trans Am Acad Ophthalmol Otolaryngol 48, 1–15 (1944).

24. Luts, H., Jansen, S., Dreschler, W. & Wouters, J. Development and normative data for the flemish/dutch matrix test (2014).

25. Brand, T. & Kollmeier, B. Efficient adaptive procedures for threshold and concurrent slope estimates for psychophysics and speech intelligibility tests. The J. Acoust. Soc. Am. 111, 2801–2810 (2002).

26. Universiteit van vlaanderen. https://www.universiteitvanvlaanderen.be/podcast. Accessed: 2022-10-20.

27. Algoet, A. Invloed van het geslacht van de spreker en luisteraar en persoonlijke appreciatie van het verhaal op de neurale tracking van de spra akomhullende. (2020).

28. Francart, T., Van Wieringen, A. & Wouters, J. Apex 3: a multi-purpose test platform for auditory psychophysical experiments. J. neuroscience methods 172, 283–293 (2008).

29. Inc., N. D. Krios. ndi. (2023).

30. Somers, B., Francart, T. & Bertrand, A. A generic EEG artifact removal algorithm based on the multi-channel Wiener filter. J. Neural Eng. 15, 036007, 10.1088/1741-2552/aaac92 (2018).

31. Søndergaard, P. & Majdak, P. The auditory modeling toolbox. In Blauert, J. (ed.) The Technology of Binaural Listening, 33–56 (Springer, Berlin, Heidelberg, 2013).

32. Biesmans, W., Das, N., Francart, T. & Bertrand, A. Auditory-inspired speech envelope extraction methods for improved eeg-based auditory attention detection in a cocktail party scenario. IEEE Transactions on Neural Syst. Rehabil. Eng. 25, 402–412 (2017).

33. Pernet, C. R. et al. Eeg-bids, an extension to the brain imaging data structure for electroencephalography. Sci. data 6, 1–5 (2019).

34. Gorgolewski, K. J. et al. The brain imaging data structure, a format for organizing and describing outputs of neuroimaging experiments. Sci. data 3, 1–9 (2016).

35. Harris, C. R. et al. Array programming with NumPy. Nat. 585, 357–362, 10.1038/s41586-020-2649-2 (2020). Number: 7825 Publisher: Nature Publishing Group.

36. Etard, O. & Reichenbach, T. Neural speech tracking in the theta and in the delta frequency band differentially encode clarity and comprehension of speech in noise. J. Neurosci. 39, 5750–5759, 10.1523/JNEUROSCI.1828-18.2019 (2019). https://www.jneurosci.org/content/39/29/5750.full.pdf.

37. Sharon, R. A., Narayanan, S., Sur, M. & Murthy, H. A. An empirical study of speech processing in the brain by analyzing the temporal syllable structure in speech-input induced eeg. In ICASSP 2019 - 2019 IEEE International Conference on Acoustics, Speech and Signal Processing (ICASSP), 4090–4094, 10.1109/ICASSP.2019.8683572 (2019).

38. Ding, N. & Simon, J. Z. Cortical entrainment to continuous speech: functional roles and interpretations. Front. Hum. Neurosci. 8 (2014).

39. Puffay, C. et al. Relating eeg to continuous speech using deep neural networks: a review. arXiv preprint arXiv:2302.01736 (2023).

40. Gramfort, A. et al. MEG and EEG data analysis with MNE-Python. Front. Neurosci. 7, 1–13, 10.3389/fnins.2013.00267 (2013).

41. Brennan Jonathan R. & Hale John T. EEG Datasets for Naturalistic Listening to “Alice in Wonderland”, https://deepblue.lib.umich.edu/data/concern/data_sets/bg257f92t (2018).

42. Vanheusden, Frederique J et al. Dataset for: Hearing aids do not alter cortical entrainment to speech at audible levels in mild-to-moderately hearing-impaired subjects, https://eprints.soton.ac.uk/438737/(2019).

